# Gene expression and DNA methylation changes in response to hypoxia in toxicant-adapted Atlantic killifish (*Fundulus heteroclitus*)

**DOI:** 10.1101/2024.11.01.620405

**Authors:** Neelakanteswar Aluru, Yaamini R. Venkataraman, Christopher S. Murray, Veronica DePascuale

## Abstract

Coastal fish populations are threatened by multiple anthropogenic impacts, including the accumulation of industrial contaminants and the increasing frequency of hypoxia. Some populations of the Atlantic killifish (*Fundulus heteroclitus*), like those in New Bedford Harbor (NBH), Massachusetts, have evolved a resistance to dioxin-like polychlorinated biphenyls (PCBs) that may influence their ability to cope with secondary stressors. To address this question, we compared hepatic gene expression and DNA methylation patterns in response to mild or severe hypoxia in killifish from NBH and Scorton Creek (SC), a reference population from a relatively pristine environment. We hypothesized that NBH fish would show altered responses to hypoxia due to trade-offs linked to toxicant resistance. Our results revealed substantial differences between populations. SC fish demonstrated a dose-dependent changes in gene expression in response to hypoxia, while NBH fish exhibited a muted transcriptional response to severe hypoxia. Interestingly, NBH fish showed significant DNA methylation changes in response to hypoxia, while SC fish did not exhibit notable epigenetic alterations. These findings suggest that toxicant-adapted killifish may face trade-offs in their molecular response to environmental stress, potentially impacting their ability to survive severe hypoxia in coastal habitats. Further research is needed to elucidate the functional implications of these epigenetic modifications and their role in adaptive stress responses.

**Summary Statement:** This study reveals how evolved resistance to toxicants in killifish may compromise their ability to respond to hypoxia, highlighting trade-offs that impact survival in stressed coastal environments.

## Introduction

Adaptation to environmental stressors is critical for survival in rapidly changing ecosystems. Understanding the physiological and molecular responses that underlie adaptive mechanisms is essential for predicting organismal sensitivity. Fish populations, particularly those inhabiting coastal waters, often face multiple environmental challenges simultaneously, which can compound the stress on these organisms. One such common stressor is hypoxia—low oxygen levels in the environment—which is increasingly documented in coastal regions due to anthropogenic activities leading excess nutrient loading, harmful algal blooms and climate change [1]. The presence and accumulation of anthropogenic chemicals in coastal ecosystems poses an additional, co-occurring threat to the health of fish populations. The ability of fish to cope with hypoxia is mediated through a range of physiological, transcriptional, and epigenetic mechanisms. However, very little is known about how populations chronically exposed to toxicants respond to secondary stressors such as hypoxia.

The Atlantic killifish (*Fundulus heteroclitus*) is one of the most ecologically important estuarine fish distributed along the East coast of the United States. Their ability to tolerate wide changes in environmental conditions, including temperature, salinity, oxygen and pH, have made them an ideal model species to investigate the biochemical, physiological and evolutionary basis of environmental adaptation [2–4]. Some populations of killifish are also valuable models for understanding the mechanisms of evolved resistance to toxicants [5]. Populations of killifish inhabiting contaminated coastal waters along the North Atlantic U.S. coast have evolved resistance to some contaminants representing major categories of aryl hydrocarbon pollutants, such as polynuclear aromatic hydrocarbons (PAHs), and halogenated aromatic hydrocarbons such as polychlorinated biphenyls (PCBs), 2,3,7,8-tetrachlorodibenzo-p-dioxin (TCDD, ‘dioxin’) and other dioxin-like compounds (DLCs) [6]. This evolved resistance involves alterations in signaling through the aryl hydrocarbon receptor (AHR), a ligand-activated transcription factor that forms a heterodimer with aryl hydrocarbon receptor translocator (ARNT), binding to dioxin-response elements to regulate the expression of target genes. While there is a great deal of understanding about the physiological and biochemical basis of adaptation to a variety of environmental conditions including toxicants in this species, very little is known about the impact of resistance to toxicants on their ability to respond to subsequent stressors such as hypoxia [7–9].

The hypoxia-inducible factor (HIF) signaling pathway is a critical cellular response mechanism that enables organisms to respond to hypoxic conditions. Under normal oxygen levels (normoxia), HIF-α subunits (mainly HIF-1α and HIF-2α) are hydroxylated by prolyl hydroxylase enzymes, marking them for degradation via the von Hippel-Lindau (VHL) ubiquitin-proteasome pathway [10, 11]. This prevents the accumulation of HIF-α under normoxia. Under hypoxic conditions, the activity of prolyl hydroxylases is inhibited due to a lack of oxygen, leading to the stabilization of HIF-α [12]. Once stabilized, HIF-α translocates into the nucleus, where it dimerizes with ARNT, also known as HIF-1β. This HIF-α/ARNT complex binds to hypoxia response elements (HREs) and regulates the transcription of target genes [13, 14]. Some of the target genes regulate processes such as angiogenesis (e.g., VEGF), erythropoiesis, glucose metabolism (e.g., GLUT1), and anaerobic metabolism (e.g., LDHA) [15]. Similar responses were observed in killifish, suggesting conserved physiological and molecular mechanisms across species [2–4, 16–20].

The AHR and HIF pathways exhibit crosstalk primarily through their shared use of the heterodimerizing partner ARNT [21]. Since ARNT is a limiting factor, competition between AHR and HIF for ARNT can influence the balance of responses to environmental toxins and hypoxic stress. This crosstalk may result in altered cellular outcomes, particularly in situations where both pathways are activated simultaneously, such as under environmental stress. One study tested this hypothesis in Atlantic killifish by exposing them to a dioxin-like PCB for three days, followed by a hypoxia challenge [22]. Prior PCB exposure disrupted the classical hypoxia response by increasing hepatic glycolytic enzyme activity, suggesting that dioxin-induced AHR activation could limit ARNT availability for the hypoxia response.

The objective of this study was to investigate the impact of evolved resistance to toxicants on response to acute hypoxia. We characterized the hepatic gene expression, and DNA methylation patterns in response to two levels of hypoxia in two distinct populations of Atlantic killifish (*Fundulus heteroclitus*). One population is from Scorton Creek, Sandwich, MA (SC), a relatively pristine environment, and is considered sensitive to environmental toxicants [23]. The other population originates from New Bedford Harbor, MA (NBH), a Superfund site heavily contaminated with dioxin-like PCBs, where the killifish have evolved resistance to toxicants. The NBH population represents a unique case study in how toxicant-adapted organisms might exhibit trade-offs in their ability to cope with additional stressors, such as hypoxia. We hypothesized that fish from NBH would exhibit altered responses to acute hypoxia compared to the sensitive Scorton Creek (SC) population. Specifically, we predicted that the NBH fish would show differential hepatic gene expression and DNA methylation patterns in response to hypoxia, potentially compromising their ability to mount an optimal response to low oxygen conditions compared to the SC population. Our results demonstrate substantial differences between the two populations in their transcriptional and epigenetic responses to hypoxia.

## Results

### Effect of hypoxia on loss of equilibrium

Neither mild nor severe hypoxia exposure had any effect on the loss of equilibrium during the 6-hour exposure period in fish from both populations. Upon initial transfer into the hypoxia chamber, fish from both populations exhibited a rapid swimming response for the first 10-15 minutes. This was followed by a noticeable reduction in swimming activity accompanied by rapid ventilation (opercular movements). By the end of the exposure period, the fish were consistently found at the bottom of the container, exhibiting slow opercular movements but there was no loss of equilibrium as evidenced by their ability to maintain position and coordinated movement.

### Gene expression changes in response to hypoxia

Strand-specific RNA sequencing of NBH and SC samples yielded an average of 17.2 million reads per sample. Of these, 83% of the reads were uniquely mapped to the genome. The summary of mapping statistics and read counts for annotated genes is provided in Supplementary Material (RNAseq_supplementaryInformation.xlsx). Principal component coordinate analysis revealed one NBH hypoxia sample to be an outlier, which was omitted from statistical analysis (Supplementary Figure S1).

#### Scorton Creek

We observed a dose-dependent effect of hypoxia on differential gene expression in SC fish. Exposure of SC fish to mild hypoxia revealed 2,241 differentially expressed genes (DEGs), with 1,170 upregulated and 1,071 downregulated. In response to severe hypoxia, 4191 DEGs were observed, with 2,221 upregulated and 1,970 downregulated. A total of 1794 DEGs were shared between the two hypoxia groups, with 980 upregulated and 814 downregulated genes (**Figure 2A**). Gene Ontology (GO) analysis of DEGs from mild and severe hypoxia treatment groups revealed overrepresentation of GO molecular function (MF) terms related to ATPase activity, RNA binding, proteosome and extracellular matrix functions. The list of top 10 overrepresented GO:MF terms among up- and downregulated DEG in SC mild and severe hypoxia groups are shown in **Figure 3**.

**Figure 1.**
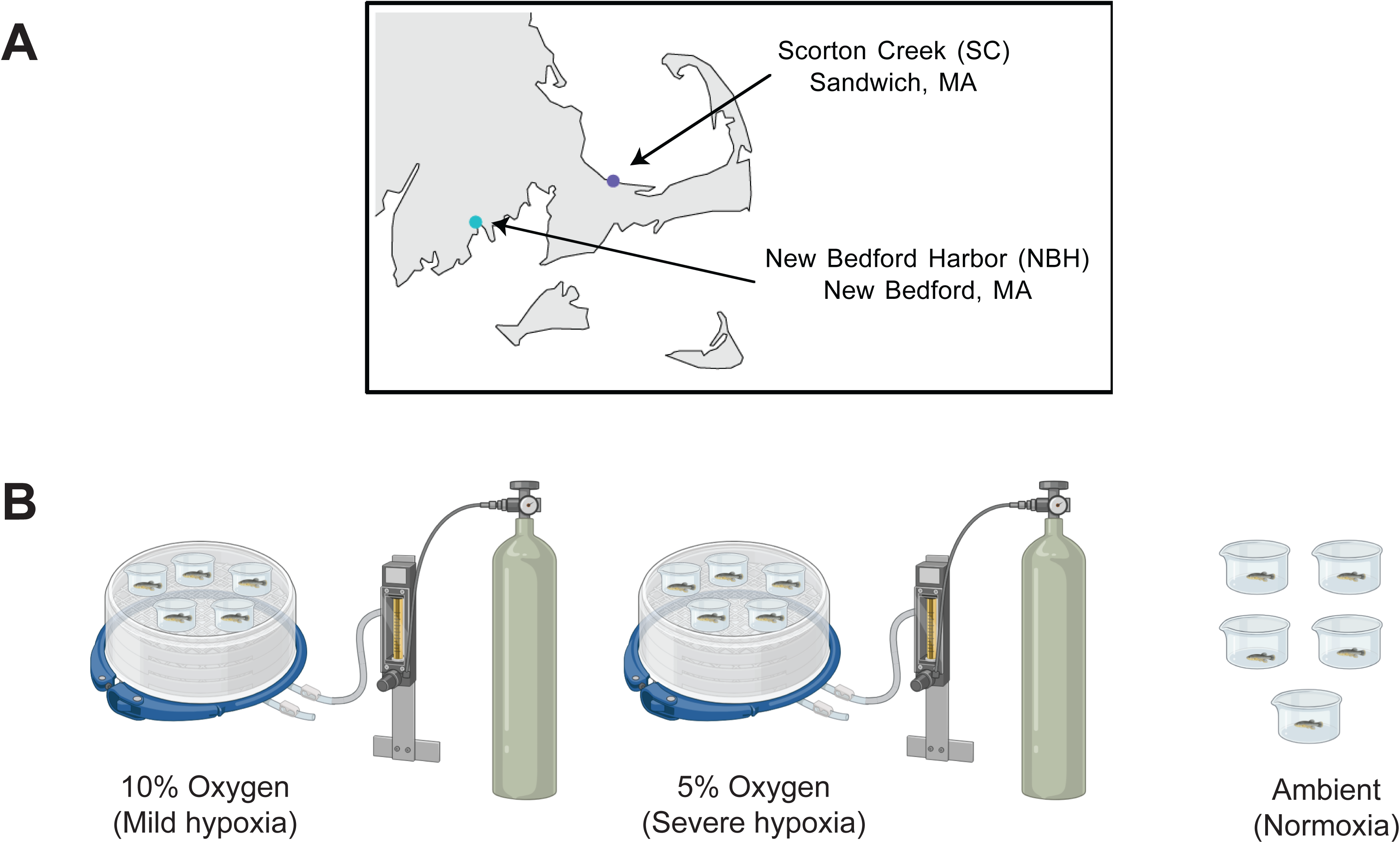
Experimental overview. **(A)** Map of Southeastern Massachusetts showing the collection sites of sensitive (Scorton Creek (SC), Sandwich, MA) and resistant (New Bedford Harbor (NBH), New Bedford, MA) Atlantic killifish. **(B)** Illustration of the experimental setup. Mild and severe hypoxia exposures were conducted by pumping oxygen containing either 5% or 10% air saturation into the chambers, respectively. Control group was maintained outside under ambient conditions.

**Figure 2.**
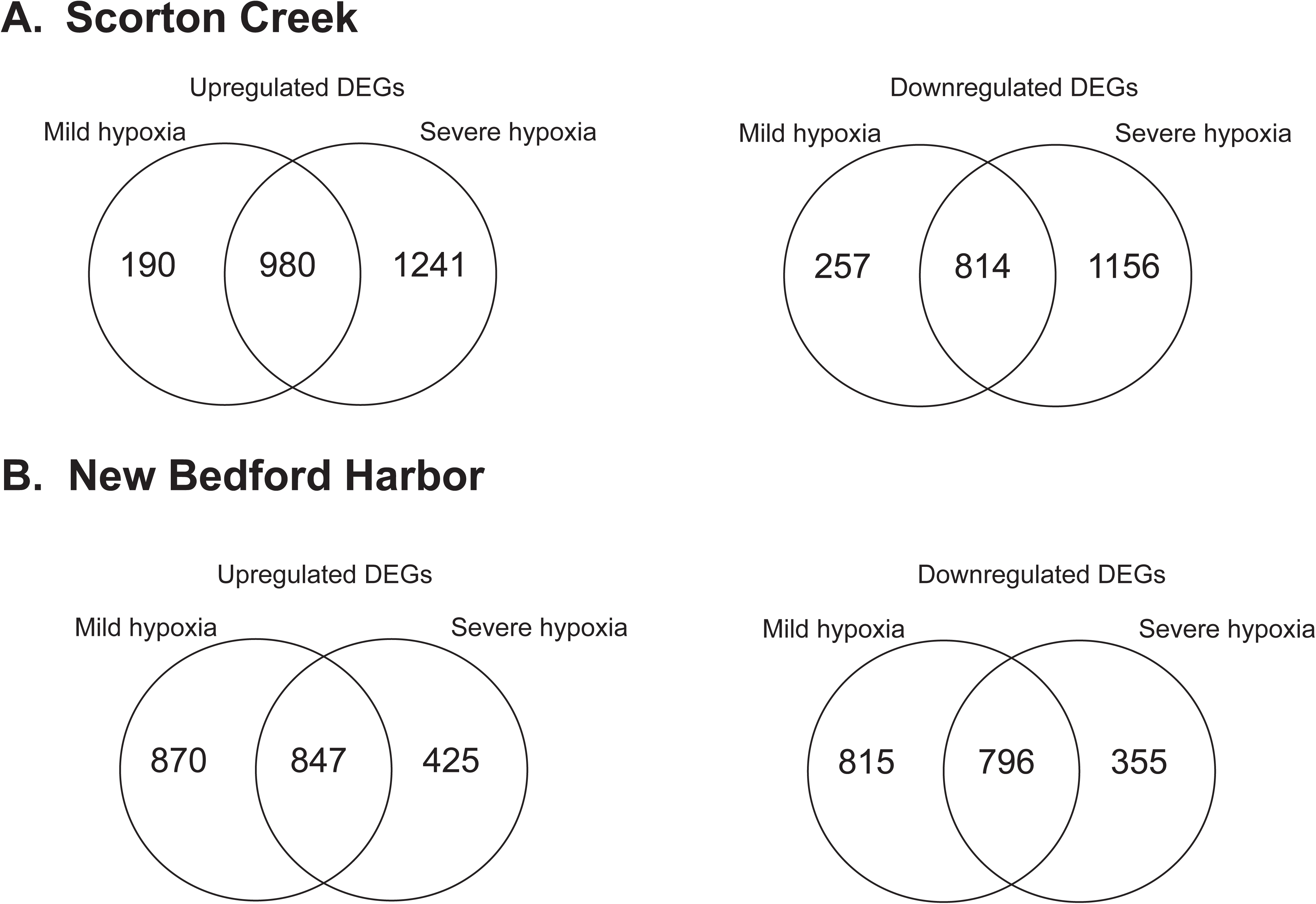
Differentially expressed genes in response to hypoxia in NBH and SC fish. Venn diagrams showing unique and common genes in response to mild and severe hypoxia in **(A)** SC fish and **(B)** NBH fish.

**Figure 3.**
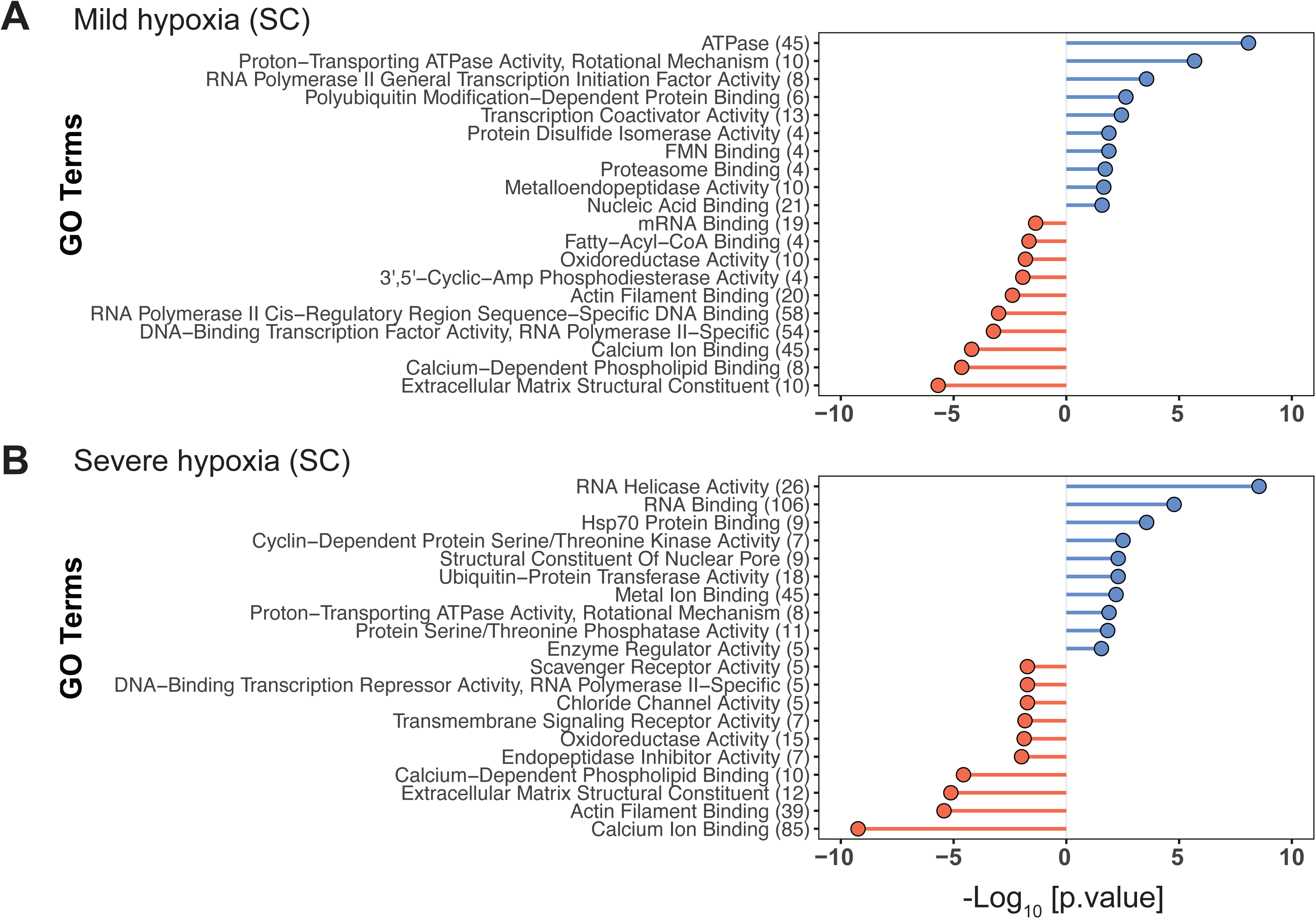
Gene Ontology (Molecular Function) terms enriched among differentially expressed genes (DEGs) in mild hypoxia **(A)** and severe hypoxia **(B)** treatment groups in SC fish. Only top 10 terms enriched among up- and down-regulated genes are shown. Entire list of GO biological process and molecular function terms are provided in the supplementary information. The numbers in the parenthesis represent the number of DEGs represented in each GO term. Detailed description of filtering of GO terms to remove redundancy is described in the materials and methods section. GO terms enriched among upregulated DEGs are in blue and those from downregulated genes are in red.

#### New Bedford Harbor

In NBH fish, exposure to mild and severe hypoxia elicited differential expression of 3,328 and 2,423 genes, respectively (**Table 1**). Among the 3328 genes differentially expressed in response to mild hypoxia, 1717 were upregulated and 1611 genes were downregulated. Whereas in response to severe hypoxia, 1,272 of the 2,423 DEGs were upregulated, and 1,151 genes were downregulated. Comparison of the upregulated DEGs from the mild and severe hypoxia groups revealed 847 genes shared between the two groups (**Figure 2B**), while a similar comparison of downregulated DEGs revealed 796 shared DEGs.

GO analysis of mild and severe hypoxia upregulated DEGs showed overrepresentation of GO:MF terms related to mRNA splicing, translation, and proteasomal degradation. The terms enriched among downregulated DEGs include specific pathways such as cell adhesion and extracellular matrix functions. The top 10 overrepresented terms among up and downregulated DEGs are shown in **Figure 4**.

**Figure 4.**
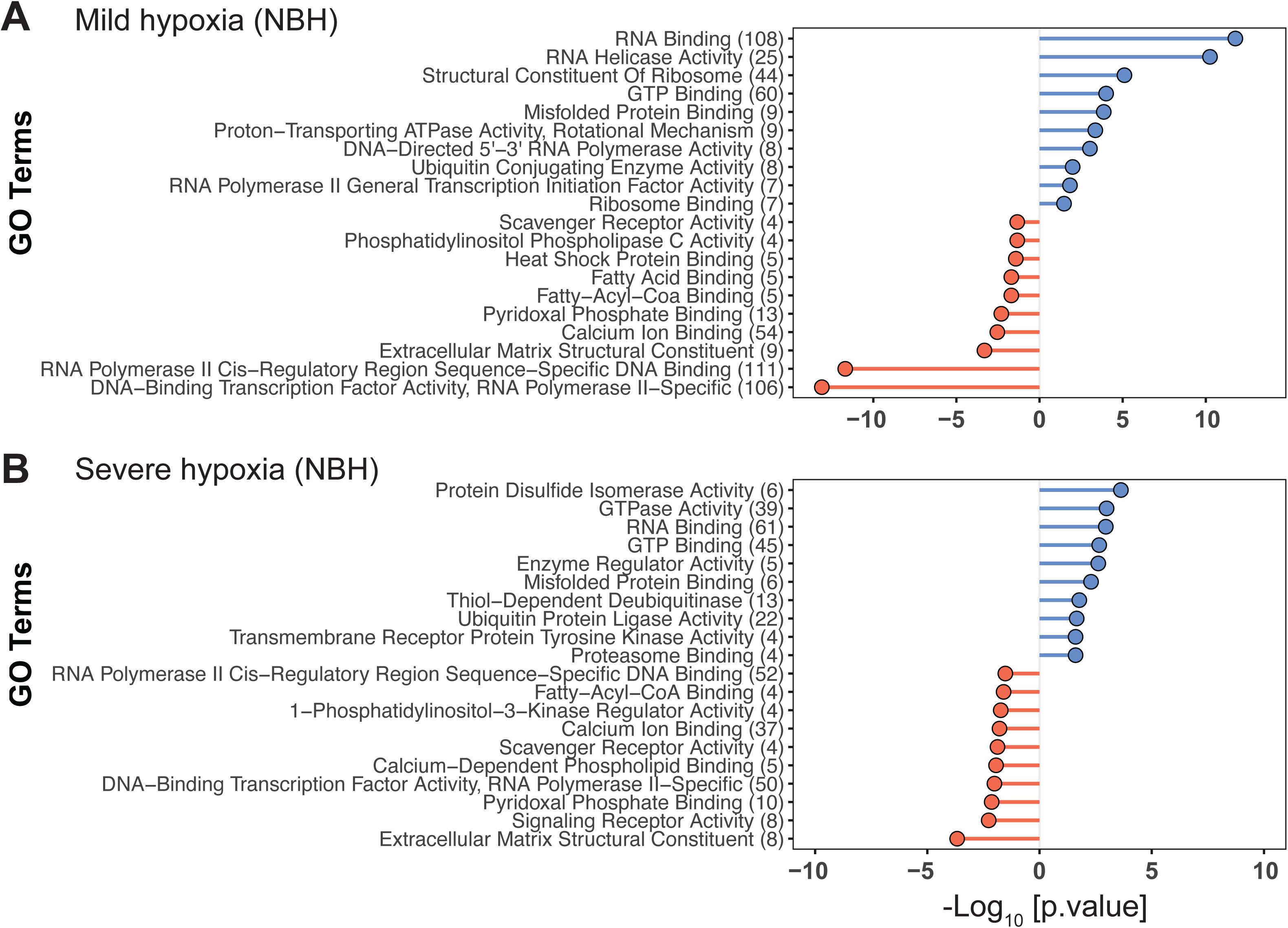
Gene Ontology (Molecular Function) terms enriched among differentially expressed genes (DEGs) in mild hypoxia **(A)** and severe hypoxia **(B)** treatment groups in NBH fish. Only top 10 terms enriched among up- and down-regulated genes are shown. Entire list of GO biological process and molecular function terms are provided in the supplementary information. The numbers in the parenthesis represent the number of DEGs represented in each GO term. Detailed description of filtering of GO terms to remove redundancy is described in the materials and methods section. GO terms enriched among upregulated DEGs are in blue and those from downregulated genes are in red.

#### Population Differences

Comparison of NBH and SC control groups revealed differential expression of 307 genes. Among them 159 and 148 are up- and downregulated in NBH, respectively, in comparison to SC. The GO terms enriched among these genes are shown in Supplementary Figure S2. Comparison of mean expression (log counts per million) of all the DEGs demonstrates that NBH fish have a more muted gene expression response to severe hypoxia in comparison to SC fish (**Figure 5**).

**Figure 5.**
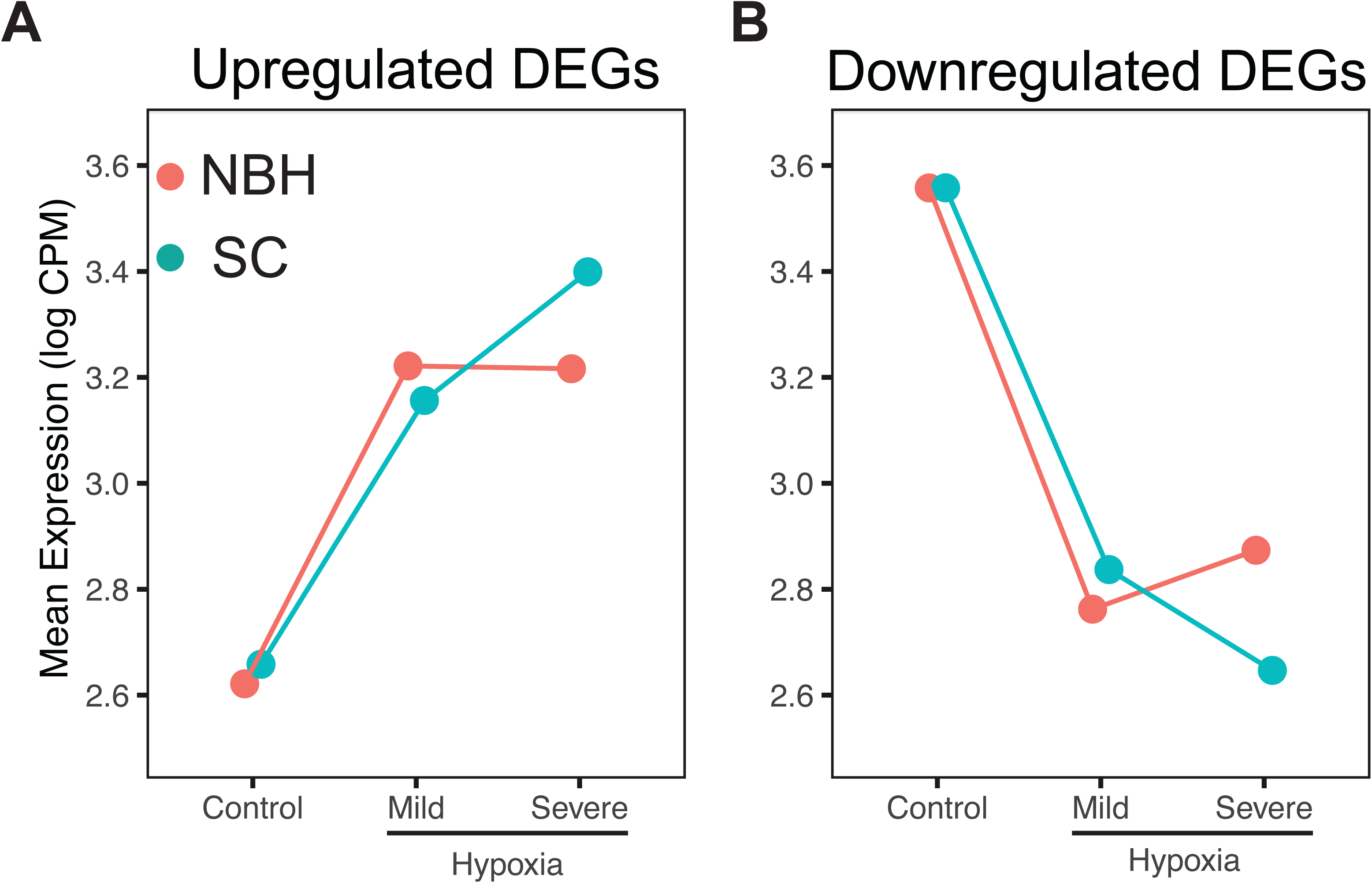
Reaction norm plots showing gene expression patterns in response to two levels of hypoxia in NBH and SC fish. Mean expression (Log counts per million (cpm)) of all the differential expressed genes in response to mild and severe hypoxia were plotted for up-**(A)** and downregulated **(B)** genes in NBH and SC fish.

We compared the two hypoxia treatment groups between NBH and SC to identify unique genes across treatments. A total of 453 upregulated and 349 downregulated genes were shared between the two populations and two hypoxia treatment groups (**Figure 6**). Heatmap representation of these genes shows magnitude of change between the two hypoxia groups in both populations (Supplementary Figure S3). Overrepresented GO terms of common up- and downregulated genes are shown in Supplementary Figure S4. These terms are similar to those observed in SC and NBH in response to hypoxia.

**Figure 6.**
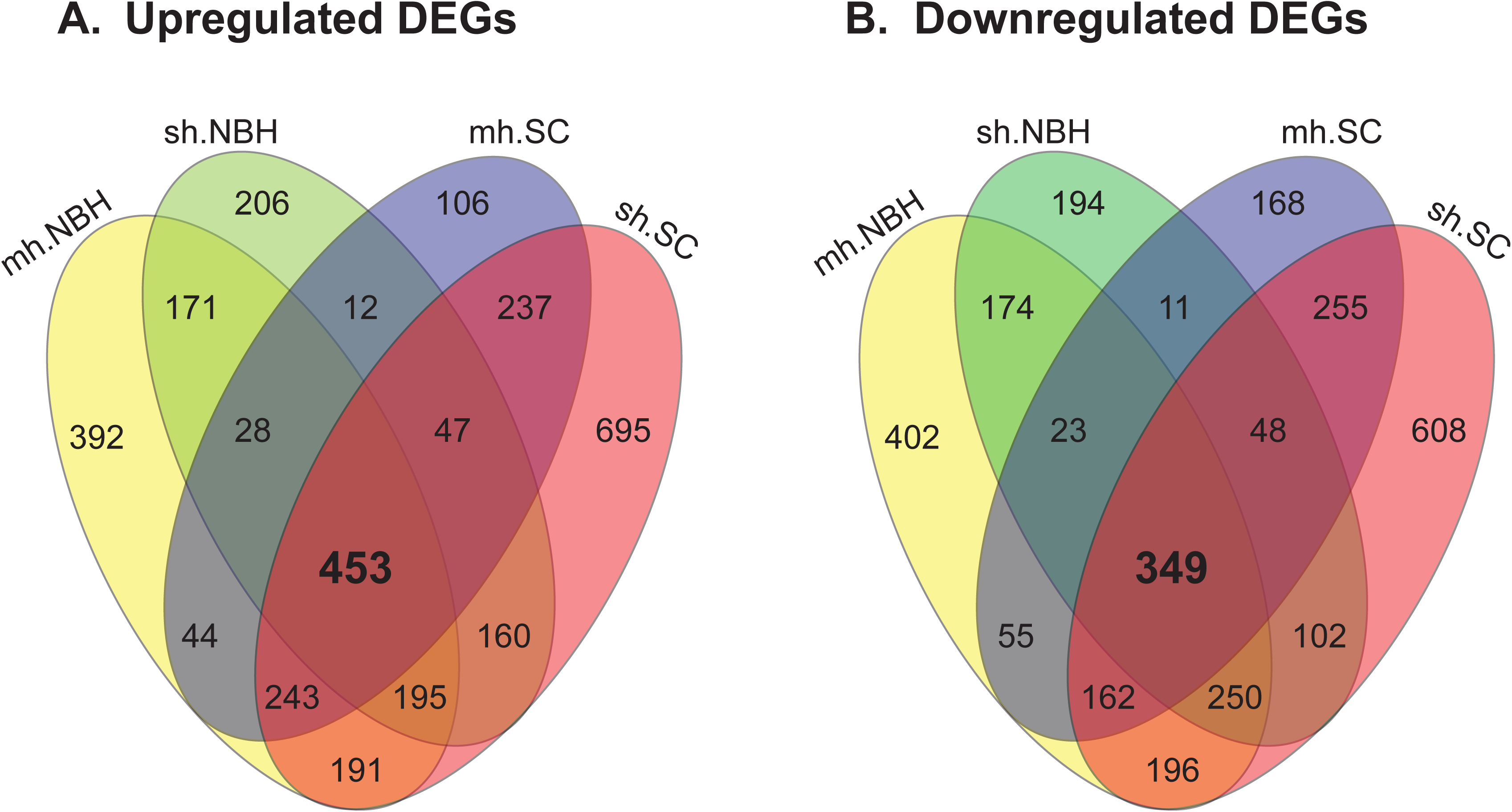
Comparison of all treatment groups. Venn diagram of upregulated (A) and downregulated (B) genes in all treatment groups revealed a core set of genes (802 DEGs) altered by hypoxia exposure irrespective of the level of hypoxia and population. GO analysis and heatmap representation of these core genes are provided in the supplementary information.

### DNA methylation changes in response to hypoxia

Reduced representation bisulfite sequencing yielded 547.1 million total paired reads (14.4 to 33.3 million paired reads per sample). Of these reads, 544.5 million (99.5%) remained after quality-trimming. From the trimmed reads, 88.6% to 94.5% total reads were aligned to the *F. heteroclitus* genome. The detailed list of mapping statistics, raw and trimmed reads per sample is provided in the supplementary information (RRBS_Supplementary_information.xlsx). A total of 439,469 CpGs (5.4% of the 8,094,243 CpGs in the *F. heteroclitus* genome) had between 10× and 500× coverage in at least one sample after CpG filtering. Among them, 148,752 CpG sites (35.2% methylated and 64.8% unmethylated) were present in all samples and were used in downstream analysis.

There was no statistically significant difference in global methylation level between NBH (28.2%) and SC liver samples (28.8%). **Figure 7** shows the mean CpG methylation density plots and total number of CpG sites in all the treatment groups. Only one DMR was identified between NBH and SC fish exposed to control conditions. This DMR is in chromosome 10 in an intron of SHISA6, a gene that encodes AMPA glutamate receptor subunit (Klassen et al., 2016). This DMR also overlaps with an annotated CpG island (identified as CpG island 77 in the genome) and is hypomethylated in NBH in comparison to SC fish.

**Figure 7.**
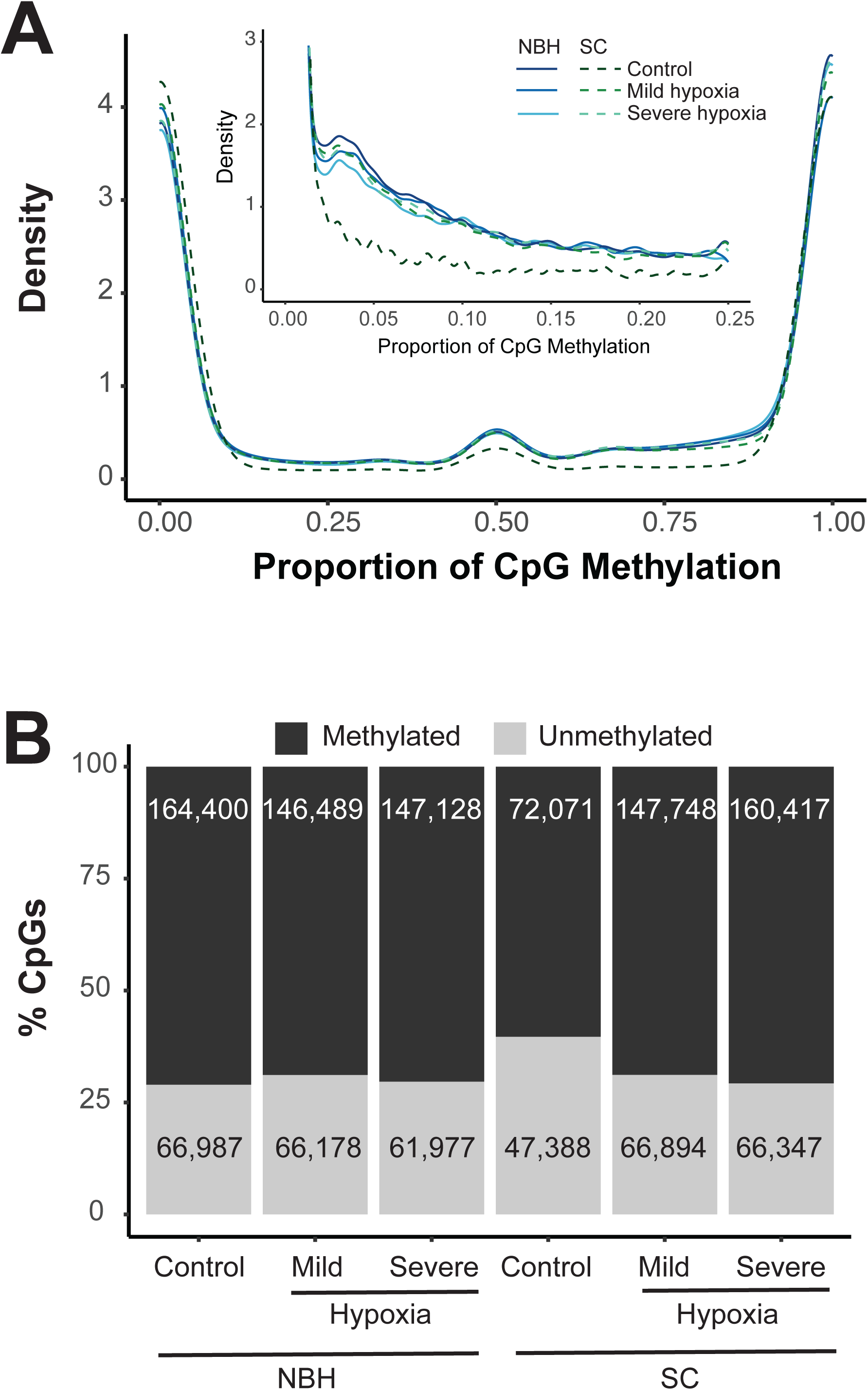
DNA methylation landscape in *F. heteroclitus*. **(A)** CpG DNA methylation density plots showing proportion of CpG methylation in different population and treatment groups. Inset shows the density plots of CpG sites with methylation levels below 25%. **(B)** Percent of methylated (>0% methylation; dark grey) and unmethylated (0% methylated; light grey) CpGs in various population and treatment groups. The average number of methylated and unmethylated CpGs are shown.

Hypoxia exposure did not substantially alter global DNA methylation levels in both populations. In SC fish, global CpG DNA methylation levels were 28.8% in the controls and mild hypoxia, and 29.2% in severe hypoxia group. No DMR were observed in response to mild or severe hypoxia in this population. In NBH fish, global DNA methylation level was 27.9% in the control group, 27.2% in mild hypoxia, and 29.7% in severe hypoxia. Comparison of control and mild hypoxia groups revealed 10 DMR, all of which were hypermethylated. Similar comparison between control and severe hypoxia revealed 59 DMRs. Among them five were hypomethylated and 54 were hypermethylated (**Figure 8**). The majority of the NBH DMR were found in CpG islands annotated in the genome. Only two DMR were shared between the mild and severe hypoxia groups, and they were hypermethylated. The genomic coordinates of these DMR are provided in supplementary information. We did not observe any significant correlation between DNA methylation in DMRs and the expression level of the associated genes (Supplemental Figures S5 and S6.)

**Figure 8.**
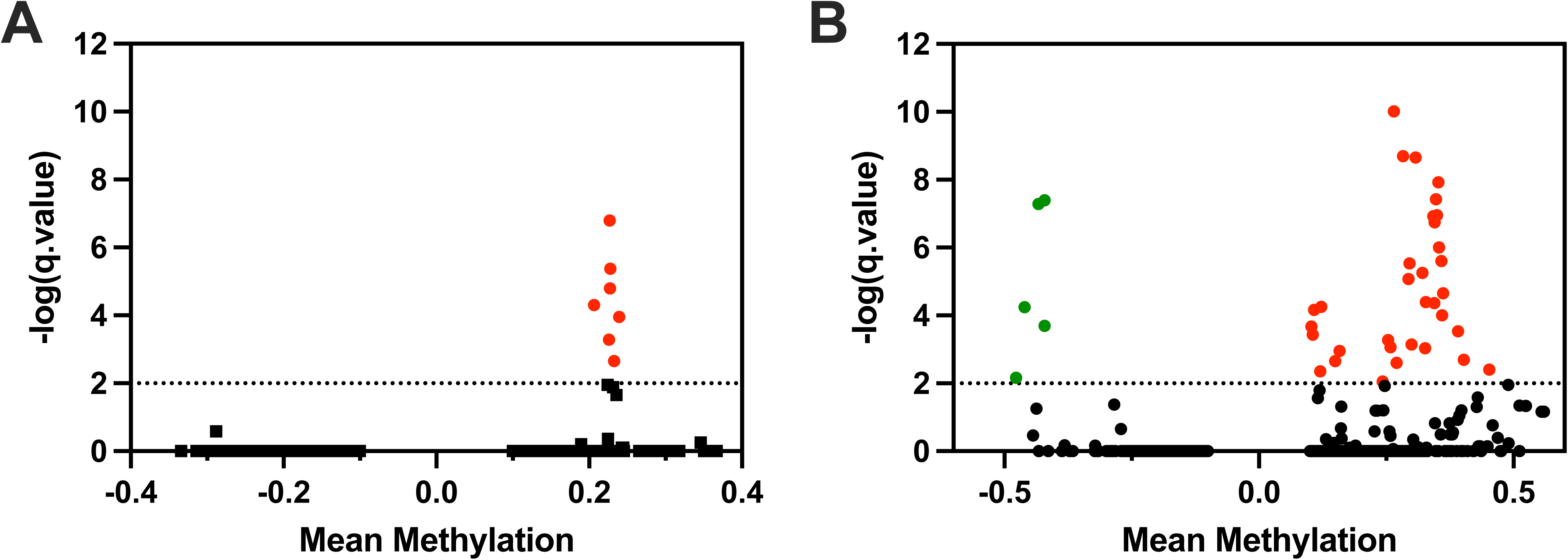
Volcano plot showing differentially methylated regions (DMRs) in response to **(A)** 10% (mild) and **(B)** 5% (severe) hypoxia exposure in New Bedford Harbor fish. Mean methylation difference (x-axis) between severe hypoxia and control group is plotted against q-value (y-axis). Each green and red spot represents a statistically significant hypo- and hypermethylated region, respectively.

## Discussion

The findings from this study demonstrate a shift in physiological responses in fish adapted to toxicants and this could compromise their capacity to respond to secondary stressors such as hypoxia. Our results show that both toxicant-sensitive (SC) and resistant killifish (NBH) respond to hypoxia but there are considerable differences in their responses as measured by gene expression and DNA methylation patterns. Atlantic killifish from SC show dose dependent changes in gene expression patterns in response to hypoxia and no changes in DNA methylation patterns, whereas the fish from NBH showed a muted gene expression response to severe hypoxia suggesting compromised ability to mount a stress response. However, NBH fish showed a modest but significant DNA methylation changes in response to hypoxia. Together, these results reveal different response mechanisms and strategies to cope with hypoxia between sensitive and resistant fish populations.

### HIF signaling in toxicant-sensitive and toxicant-resistant populations

Transcriptional responses to hypoxia are well-documented in fish species [24–28], including Atlantic killifish [29–31]. Our findings demonstrate that both SC and NBH killifish exhibit responses to hypoxia, though their reactions differ significantly. The hypoxia-inducible transcription factors (HIF-1α, HIF-2α, HIF-3α) play a major role in the transcriptional activation of the hypoxic response [32]. The expression of these genes under hypoxic conditions depends on the intensity and duration of hypoxia exposure. Interestingly, we did not observe any changes in the expression of HIF1α and HIF2α genes in response to mild or severe hypoxia in either of the populations. However, we observed an increased expression of HIF3α only in NBH fish (mild hypoxia – log FC 1.75, FDR 7.45E-04; severe hypoxia – log FC 1.54, FDR 4.07E-03). A previous study in a closely related species, *Fundulus grandis*, showed no significant differences in mRNA expression of any of these HIF genes in response to hypoxia (1 mg oxygen/L) after 6 and 24 hours post-exposure [33]. However, studies in mammalian cell culture systems have shown differences in temporal profiles of HIF genes in response to hypoxia. In human endothelial cells, HIF-1α expression is maximal after 4 hours of hypoxia and then is dramatically reduced by 8 hours [34]. In contrast, HIF-2α is maximal at 8 hours and remains elevated up to 24 hours [34–36]. Unlike HIF-1α and -2α, moderate hypoxia exposure induced HIF3α mRNA expression within 2 hours [35]. However, very little is known about the functional role of HIF3α in hypoxia responses. The upregulation of HIF3α in response to hypoxia only in NBH fish suggests that adaptation to contaminated environment may have altered its regulation, with potential implications for the hypoxia response. Further studies are needed to understand the roles of all three HIFs under different hypoxic conditions and at different life stages.

Another set of HIF pathway genes that were differentially expressed are the prolyl hydroxylase domain-containing proteins (PHD) [10]. There are three 2-oxoglutarate-dependent PHD proteins (PHD1, PHD2, and PHD3, encoded by EGLN2, EGLN1, and EGLN3, respectively) that are involved in HIFα ubiquitination and proteasomal degradation under normoxic conditions [37]. Hypoxia has been shown to induce the expression of EGLN1 and -3 mRNAs, but not EGLN2, in several cell types [12]. In most cases, the induction of EGLN3 mRNA is much more prominent than that of EGLN1. We observed a several-fold increase in the *egln3* expression (SC: log FC 8.77, FDR 1.82E-06; NBH: log FC 3.51, FDR 0.028) and modest but significant *elgn1a* upregulation in response to severe hypoxia in both populations. We also observed upregulation of *egln2* (logFC 1.33, FDR 9.59E-04) in response to severe hypoxia in SC fish, while it was not significant in NBH fish. Although very little is known about the role of *egln2* in oxygen sensing and hypoxia tolerance, the differences in expression between the two populations are intriguing. It has been suggested that increase in EGLN activity during hypoxia acts as a regulatory feedback loop for fast elimination of HIF after reoxygenation [38].

In addition to the direct regulation of HIF transcription by hypoxia, multiple signaling pathways have been shown to play a role in the regulation of HIF gene expression in a variety of model systems. These include the PI3K-mTOR, interleukin-6 (IL-6), ERK, and MAPK signaling pathways [15]. We observed differential expression of several genes associated with these signaling pathways only in NBH fish, suggesting a role in adaptation to toxicants. Despite the lack of an increase in HIF-1α expression, the differential expression of these signaling pathways suggests that NBH fish utilize cross-talk between multiple signaling pathways to cope with hypoxia in a way that appears to be distinct from SC fish.

### Transcriptional responses to hypoxia

While responses to hypoxia are well-documented across various organisms, encompassing metabolic changes, angiogenesis, and vascularization [3, 4, 15, 16], our results, based on a 6-hour hypoxia exposure, revealed gene expression changes that were more focused on transcription, translation, and cell cycle-related genes, rather than the classical hypoxia responses. This contrasts with studies in *in vitro* mammalian systems, where hypoxia target genes are differentially expressed in shorter time scales [39]. The lack of similar responses in our study could be due to differences in the timescales of transcription and translation between fish maintained at 20°C and mammalian cells held at 37°C [39, 40]. SC fish exhibited dose-dependent changes in gene expression, indicating a gradual and proportional response to varying levels of hypoxia. In contrast, NBH fish demonstrated drastic gene expression changes in response to mild hypoxia, but a muted response to severe hypoxia. This pattern suggests that NBH fish have a lower tolerance threshold for hypoxic conditions and are unable to induce a robust transcriptional response to severe hypoxia. This could be potentially due to higher energetic costs for survival in contaminated environment leading to compromised ability to respond to additional stressors [7, 8].

In SC fish, mild hypoxia exposure caused an overrepresentation of genes associated with efflux transporters, transcription factor activity, mRNA and nucleic acid binding, and proteasomal functions. All these functions have been previously shown to be altered by hypoxia [11, 41–44]. For instance, efflux pumps belonging to the ATP-binding cassette (ABC) superfamily of membrane transporters are known to play a significant role in cellular protection against oxidative stress [45]. Several genes within the ABC family of efflux transporters are expressed in the liver and are involved in the transport of glutathione and glucuronide conjugates [46–48]. These findings suggest that mild hypoxia increases reactive oxygen species (ROS), with glutathione playing a crucial role in neutralizing free radicals and protecting the liver. The glutathione conjugates are subsequently eliminated from hepatocytes by the efflux pumps, contributing to cellular detoxification [48]. Additionally, mild hypoxia caused the upregulation of genes associated with RNA polymerase II (RNAPII) general transcription factor activity and the downregulation of cis-regulatory sequence-specific DNA binding RNAPII activity. This is not surprising, given the evidence that hypoxia affects the transcription of hundreds of genes [13]. Under normoxic conditions, almost all HIF target genes display an open chromatin structure and harbor transcriptionally active but paused RNA polymerase II [49]. Changes in the expression of RNAPII activity genes in response to hypoxia suggests the release of paused Pol II into productive RNA synthesis by recruiting various coactivators, repressors, and chromatin remodelers, resulting in either the activation or inhibition of transcription of target genes [49].

In response to severe hypoxia, SC fish showed an overrepresentation of genes associated with RNA helicases. They play an important role in cellular RNA metabolism, including transcription, pre-mRNA splicing, RNA export, storage, decay, and translation. Recently, RNA helicases have been shown to be involved in many biological processes, including DNA damage repair, cellular stress response, hypoxia and antiviral defense [50, 51]. Another major overrepresented group of genes in response to severe hypoxia in SC fish are the RNA binding proteins (RBPs) – important players in mRNA turnover (decay and stabilization) and translation [52]. RNA helicases and RBPs have been shown to be essential for the adaptive cellular response to hypoxia [50, 53]. While there is very little understanding on the role of RBPs in environmental model species such as killifish, the upregulation of these genes in response to hypoxia suggests highly conserved cellular mechanisms.

Interestingly, the genes and pathways that were upregulated in SC fish under severe hypoxia were similarly upregulated in NBH fish exposed to mild hypoxia. This suggests that the toxicant-adapted NBH fish are more sensitive to mild hypoxia, undergoing more drastic transcriptional changes. The upregulation of RNA helicases and RBPs indicates post-transcriptional regulation of pre-existing RNAs [53], suggesting that NBH fish may reduce transcription under stress. Indeed, this was observed under severe hypoxia in NBH fish, where the number of DEGs was lower compared to SC fish, indicating a drastic reduction in transcription.

Hypoxia exposure downregulated genes related to calcium signaling, oxidoreductase activity, and extracellular matrix (ECM) modeling proteins. These pathways were altered in both hypoxia treatments and across both populations, suggesting they are vital for cellular response to hypoxia. It is well-established that hypoxia impairs mitochondrial respiration and ATP synthesis, leading to an increased production of reactive oxygen species (ROS) and calcium release from the ER into the cytosol [15]. The resulting elevated cytosolic calcium levels cause increased calcium uptake into mitochondria and mitochondrial calcium overload, which in turn leads to mitochondrial depolarization and the initiation of cell death [54]. The decreased expression of genes associated with calcium signaling suggests an adaptation toward hypoxia tolerance. Similarly, hypoxia involves a switch from oxidative phosphorylation to glycolysis, resulting in increased production of NADH and an imbalance in the NAD+ and NADH ratio, causing altered redox potential [14, 55]. These conditions favor the overexpression of many redox enzymes, such as cytochrome P450 reductase and nitroreductases. Even though these genes were not altered, genes that are dependent on NAD+ or NADH and play critical roles in metabolism were downregulated. They include *agmo* (alkylglycerol monooxygenase), *sc5d* (sterol-C5-desaturase), *hsd11b2* (11-β-hydroxysteroid dehydrogenase 2), *bdh1* (d-beta-hydroxybutyrate dehydrogenase), and *foxred1* (FAD-dependent oxidoreductase domain-containing 1) [14]. It remains to be determined whether the downregulation of these genes under hypoxia is adaptive or maladaptive.

Another significant group of genes that are commonly downregulated in response to hypoxia are ECM components genes. The ECM is a complex network of proteins and other molecules that provide structural and biochemical support to surrounding cells [56]. Its composition and function are crucial for tissue integrity, cell behavior, and overall organism health [57]. Hypoxia has been shown to cause significant effects on the ECM in various aquatic species [58–61]. In addition, there is growing evidence suggesting the role of hypoxia and HIFs in reprogramming cancer cells by regulating extracellular matrix (ECM) deposition, remodeling and degradation, thereby promoting cancer metastasis [62]. We observed downregulation genes associated with collagen synthesis, which are essential for maintaining the structural integrity of tissues. This could lead to weakened tissue architecture or cause tissue remodeling such as changes in vascularization, which can be adaptive to survive under hypoxic conditions. The molecular mechanisms associated with these changes could be either by direct regulation of ECM pathway genes by HIF proteins or indirectly by the induction of oxidative stress [57, 63]. Overall, the effects of hypoxia on the ECM in environmental species such as killifish could be critical, as they frequently encounter hypoxic conditions under diel cycle and ECM-related adaptations are critical for survival.

### Epigenetic changes in responses to hypoxia

Epigenetic effects, particularly DNA methylation changes in response to hypoxia, are well documented across a variety of fish species [64–68]. Hypoxia has been shown to cause both global and gene-specific alterations in DNA methylation patterns, which can influence gene expression, development, and stress response pathways. For instance, in species like zebrafish (*Danio rerio*) and Atlantic salmon (*Salmo salar*), hypoxic conditions have been associated with both hypermethylation and hypomethylation of key regulatory genes involved in metabolic adaptation and oxidative stress responses [64, 68]. These epigenetic changes are hypothesized to help fish cope with hypoxia by modulating critical pathways for survival, growth, and development.

To our knowledge, this is the first study where genome-wide DNA methylation patterns are profiled in Atlantic killifish. In this study, we hypothesized that resistant and sensitive populations of Atlantic killifish would exhibit distinct hepatic DNA methylation patterns, possibly reflecting their differential tolerance to dioxin-like PCBs. However, we did not observe distinct population differences in DNA methylation (NBH vs SC control groups), suggesting that parental exposure to dioxin-like PCBs did not have a detectable effect on DNA methylation in the offspring held under normoxic conditions.

Hypoxia exposure induced DNA methylation changes that were markedly different in the two populations. In the sensitive SC population, no differentially methylated regions (DMRs) were identified in response to either mild or severe hypoxia. In contrast, hypoxia resulted in significant changes in DNA methylation in the toxicant-resistant NBH population. Most of these DMRs are located in intronic regions and were enriched within CpG islands, which are known to play a critical role in regulating gene expression [69]. These findings suggest that the NBH population may have a more plastic epigenetic response to hypoxia compared to the SC population, which could reflect underlying differences in their adaptive capacities. While further research is needed to elucidate the functional consequences of these methylation changes, this study provides new insights into the epigenetic mechanisms by which fish populations respond to environmental stressors like hypoxia.

The majority of hypoxia induced DMRs are hypermethylated supporting previous observations that hypoxia cause DNA hypermethylation [70–72]. Associating DMRs with genes revealed that some of the DMRs are related to genes involved in sialylation, vascularization and development. Four DMRs were associated with the gene *St6galnac3*, which is involved in the sialylation pathway. Sialylation refers to the addition of sialic acid units to oligosaccharides and glycoproteins [73]. Sialic acids moieties act as bridging molecules, facilitating communication between cells and the extracellular matrix. Additionally, one DMR was associated with *beta-chimaerin*, a protein linked to vascularization [74]. Hypoxia has been shown to influence both sialylation and vascularization processes, particularly in cancer models [74, 75], suggesting that similar mechanisms may be involved in the hypoxic responses observed in this study. However, we did not observe any significant correlation between these DNA methylation changes and expression of the associated genes. None of the 58 genes associated with DMRs in NBH fish were differentially expressed suggesting a temporal lag in DNA methylation changes and gene expression. In addition, several studies have shown that majority of the DNA methylation changes are not correlated with gene expression [76–78]. This suggests that epigenetic regulation of gene expression is multilayered with many levels of control, involving DNA methylation, histone modifications and chromatin organization.

While differential methylation did not correlate with gene expression changes in NBH, we observed differential expression of several chromatin modifier genes, particularly histone lysine demethylases (KDMs), in response to hypoxia in both fish populations. This is expected, as KDMs belong to the family of 2-oxoglutarate-dependent dioxygenases, which function as oxygen sensors [13]. The differentially expressed KDM genes include *kdm1aa*, *kdm2aa*, *kdm2ab*, *kdm2ba*, *kdm3b*, *kdm4b*, *kdm5ba*, *kdm5bb*, *kdm5c*, *kdm6a*, *kdm6ba*, and *kdm7aa*. These genes have been shown to be directly regulated by hypoxia-inducible factors (HIFs), linking their expression to the cellular response to hypoxic stress [79, 80]. Given the differential expression of histone lysine demethylases in response to hypoxia, it would be intriguing to investigate the role of chromatin modifiers in the adaptation to hypoxia in environmental species, as they may be key regulators of gene expression in low-oxygen environments.

### Conclusions

This study highlights the complex and distinct physiological and epigenetic responses of Atlantic killifish populations adapted to toxicants when exposed to hypoxia. Our findings suggest that the capacity to respond to secondary stressors such as hypoxia may be altered in populations adapted to environmental contaminants. As expected, the toxicant-sensitive SC fish displayed a dose-dependent response to hypoxia exposure. However, the toxicant-resistant NBH fish, exhibited muted transcriptional responses but a more pronounced DNA methylation response to severe hypoxia, suggesting different molecular mechanisms in this population. Importantly, the differential DNA methylation patterns in response to hypoxia between the two populations indicate differences in epigenetic plasticity, which needs further investigation. Overall, this research provides valuable insights into the diverse molecular mechanisms by which fish populations, with different environmental histories, respond to hypoxia and highlights the need for further exploration of epigenetic and chromatin-level responses in the context of environmental adaptation.

## Materials and Methods

### Experimental fish

The animal husbandry and experimental procedures used in this study were approved by the Animal Care and Use Committee of the Woods Hole Oceanographic Institution. Mature adult male and female killifish from Scorton Creek (SC; Sandwich, MA) and New Bedford Harbor (NBH; New Bedford, MA) were collected using minnow traps, as described previously [81]. Fish were maintained in the Redfield Laboratory (WHOI) with continuous flow-through seawater (SW) at 18–20°C, saturated dissolved oxygen (21% oxygen saturation or 7.21 mg O_2_ L^-1^), and 14h:10h light/dark photoperiod conditions.

F1 generation of embryos from SC and NBH were obtained by *In vitro* fertilization following established protocols (Karchner et al., 1999). Briefly, 4-5 female fish from SC or NBH (15.5 ± 1.2 g mean wet mass) were lightly anesthetized with Tricaine (MS222; buffered with sodium bicarbonate, Sigma-Aldrich, St. Louis, MO, USA) and oocytes were obtained for *in vitro* fertilization by gently squeezing the abdomen. Oocytes were collected in glass petri dishes with filtered SW (30 parts per thousand; ppt). Milt was obtained by euthanizing 2-3 mature males (13.5 ± 1.1 g mean wet mass) from the same population in MS222, dissecting out the gonads, and chopping them with a scalpel blade in seawater. A few drops of milt were added to the oocytes for fertilization. Approximately 20 minutes after the addition of milt, embryos were rinsed with filtered SW to remove any excess sperm. Fertilized embryos were reared at 23°C under 14h:10h light/dark photoperiod conditions until hatching. Larvae were raised in 2-gallon aquarium tanks in aerated seawater for six months. During larval rearing, fish were fed brine shrimp daily and water was exchanged every 2-3 weeks. Oxygen concentration was measured in the tanks once every 2-3 days and oxygen saturation was above 20% throughout the rearing period.

### Hypoxia exposure

Six-month-old killifish juveniles from SC (149 ± 43 mg mean wet mass) and NBH (145 ± 39 mg mean wet mass) were exposed to either mild (10% oxygen saturation, 3.46 mg O_2_ L^-1^; n = 5 per population) or severe hypoxia (5% oxygen saturation, 1.72 mg O_2_ L^-1^ ; n = 5 per population) for 6 hours. These two hypoxia levels were chosen based on preliminary experiments with the same cohort of fish, where the loss of equilibrium (LOE) was assessed under 1% and 5% oxygen saturation in both NBH and SC fish. Six hours of exposure to 5% oxygen saturation did not cause LOE in fish from either population (n = 5 individual fish per population), whereas 1% oxygen saturation caused LOE within 6 hours in fish from both populations.

Hypoxia exposure set up includes pyrex glass dishes (270 mL volume) equipped with oxygen sensor spots (PreSens Precision Sensing GmbH, Germany) placed inside hypoxia chambers (STEMCELL Technologies Inc.) with pre-mixed air set to 5 or 10% oxygen pumped into the chambers continuously. A control group (normoxia, 20.9% oxygen saturation; n = 5 per population) was maintained on the benchtop (**Figure 1**). Prior to introducing the fish to hypoxia, 250 mL filtered seawater was added to the pyrex glass dishes and was allowed to equilibrate overnight to ensure that the water has reached the respective treatment conditions. Oxygen levels in individual beakers were checked prior to introducing the fish using a FireString oxygen sensor (PyroScience, Germany) and were found to be at the treatment conditions in each of the individual dishes. At the start of the experiment, individual fish were quickly introduced into the pyrex dishes and chambers closed quickly. Fish were maintained at treatment conditions for 6 hours. At the end of the exposure period, oxygen levels were measured.

### Isolation of total RNA and genomic DNA from liver samples

Simultaneous isolation of genomic DNA and total RNA from liver tissues was performed using the ZR-Duet DNA/RNA Mini Prep kit (Zymo Research, California). RNA was treated with DNase during the isolation process. DNA and RNA were quantified using the Nanodrop Spectrophotometer. The quality of DNA and RNA was checked using the Agilent 4200 and 2200 Tape Station systems, respectively. The DNA and RNA integrity numbers of all samples were between 9 and 10.

### RNA sequencing

Libraries were constructed using Illumina stranded library preparation kit following manufacturer’s protocol. Single end 50bp reads were sequenced using Illumina HiSeq2500 platform. RNA sequencing library construction and sequencing were done at the Tufts university core facility.

### Reduced Representation Bisulfite Sequencing (RRBS)

Library preparation was performed using the Premium RRBS kit (Diagenode). In brief, 100 ng DNA from each sample were enzymatically digested by the restriction enzyme MspI at 37°C for 12 hours. Following ends preparation, a different set of adaptors was added to each sample and adaptor ligation was performed by the addition of ligase. Size selection of adaptor-ligated DNA fragments was performed by Agencourt AMPure XP beads (Beckman Coulter) and the DNA was eluted in Resuspension buffer. Part of the eluted sample was subjected to qPCR using 2X KAPA HiFi HotStart ReadyMix (Kapa Biosystems) for quantification and subsequent pooling per 9 samples. The pooling was performed according to two parameters: the Ct value and the adaptor ID of each sample. The pooling was followed by a cleanup with AMPure XP beads to reduce the volumes. Bisulfite treatment was performed, and bisulfite-converted DNA was eluted twice in BS Elution buffer. Part of the bisulfite converted library was used in qPCR for the determination of the optimal cycle number for the enrichment PCR. 2X MethylTaq Plus Master Mix was used for the amplification PCR and a last cleanup with AMPure XP beads followed. PCR product was run on an 2% agarose gel to remove adaptor dimers. The quality of the final libraries was checked on an Agilent 2100 High Sensitivity DNA chip. The concentration was determined by performing qPCR on the samples using a dilution of PhiX index3 as standard. Paired end 50bp reads were sequenced on an Illumina HiSeq4000 platform by a commercial facility (NXT-Dx, Ghent, Belgium).

### Genome Information and Feature Tracks

The *F. heteroclitus* genome (https://www.ncbi.nlm.nih.gov/datasets/genome/GCF_011125445.2/) was used for all analyses. Genome feature information was pulled directly from the genome to generate gene, coding sequence (CDS), exon, and lncRNA genome feature tracks using Gnomom, RefSeq, cmsearch, and tRNAscan-SE annotations. Chromosome name and length information was also extracted from the genome to generate additional feature tracks using bedtools v2.31.1 [82]. A non-coding sequence track was created by using the complement of the CDS track (`complementBed`). Similarly, the intergenic genome feature track was created using the complement of the gene track. The intersection (`intersectBed`) between the non-coding sequence and gene tracks were used to create an intron track. All genome feature tracks are available in an Open Science Framework repository (doi.org/10.17605/OSF.IO/NZRA8).

### RNA sequencing and analysis

Raw data files were assessed for quality using FastQC Version 0.11.9 [83] prior to preprocessing. Preprocessing was done by trimming the adaptor sequences using Trimmomatic (Version 0.25) and removing any reads with low sequence quality (Phred score < 20) [84]. Trimmed sequence reads were mapped to the *F. heteroclitus* genome using the STAR aligner v.2.6.1d [85]. The number of reads mapped to annotated regions of the genome was obtained using HTSeq-count v.0.11.1 [86]. Statistical analysis was conducted using edgeR v.3.40.2, a Bioconductor package [87]. Transcripts from all samples were compiled into a DGEList, and lowly expressed transcripts were filtered out using the filterByExpr function. Sample ordination was visualized using multidimensional scaling analysis with the ape v5.8 package [88], revealing an outlying sample that was removed from subsequent analysis (Fig. S1). We used the quasi-likelihood model in edgeR (glmQLFTest) to perform differential gene expression analysis. Only genes with false discovery rate (FDR) of <5% were considered to be differentially expressed. Raw data has been deposited in gene expression omnibus (Accession number GSE278569).

Functional annotation of DEGs was done using gene ontology (GO) biological process (GO:BP) and molecular function (GO:MF) terms. Identification of overrepresented GO terms (*p* value < 0.05) among sets of DEGs was done using the enricher function in ClusterProfiler v.4.6.2 [89]. The background gene list included all expressed transcripts in the filtered DGEList. Similar GO:BP or GO:MF terms were clustered based on the frequency of shared genes using the R package *rrvgo* 1.16.0 [90]. Representative parent terms from each cluster were chosen based on the lowest *p value*.

### DNA methylation profiling by Reduced Representation Bisulfite Sequencing (RRBS)

The Bisulfite Analysis Toolkit (BAT) [91] was used for RRBS analysis. Prior to analysis, raw data was quality trimmed with TrimGalore! v.0.6.6 [92]. Trimming was performed on non-directional (--non_directional) paired-end reads (--paired). An additional 2 bp were trimmed from the 3’ end of the first read and 5’ end of the second read (--rrbs). Sequence quality was assessed with FastQC v0.11.9 [83] and MultiQC v1.11 [93] after trimming.

Trimmed paired reads were aligned to the genome using BAT_mapping module specifying non-directional input (-F 2). Mapping statistics and methylation calling was done using BAT_mapping_stat and BAT_calling modules, respectively. CpG methylation data was filtered to retain only a minimum 10 and maximum 500 reads per sample (--MD_min 10, –MD_max 500 -- CG). The data were sorted using bedGraphs (sortBed v2.29.1; [82]) and merged into treatment-specific groups (BAT_summarize). Within each group, one sample was allowed to have missing data for a CpG locus (--mis1 1, --mis2 1). If data were missing for more than one sample at a particular CpG, it was not included in the downstream analysis. Chromosome lengths were specified (--cs) for merging methylation information accurately. BAT_overview was used to obtain average methylation rate per sample in each group, hierarchical clustering of sample methylation rates, distribution of CpG methylation, comparison of methylation rate between groups for common loci, and differences in mean methylation rate between groups. Differentially methylated regions (DMR) — defined as regions with at least 10 CpGs, a minimum methylation rate difference of 0.1, and q-value < 0.05 — were identified for each comparison using BAT_DMRcalling module. The closest genome feature to each DMR was characterized using closestBed.

## Supporting information

RRBS_supplemental_information

RNAseq_supplemental_information

## Acknowledgements

This work is partly supported by The Richard B. Sellars Endowed Research Fund (WHOI Independent study award) and National Institutes of Health R01 ES024915 to NA. VD was supported by WHOI Summer Student Fellowship. YRV (NSF-PDF #2209018) and CSM (NSF-PDF #2126533) are supported by NSF PRFB and OCE postdoctoral fellowships, respectively.

