## Supplemental Information for "Gene expression and DNA methylation changes in response to hypoxia in toxicant-adapted Atlantic killifish (*Fundulus heteroclitus*)"

**Figure S1.**

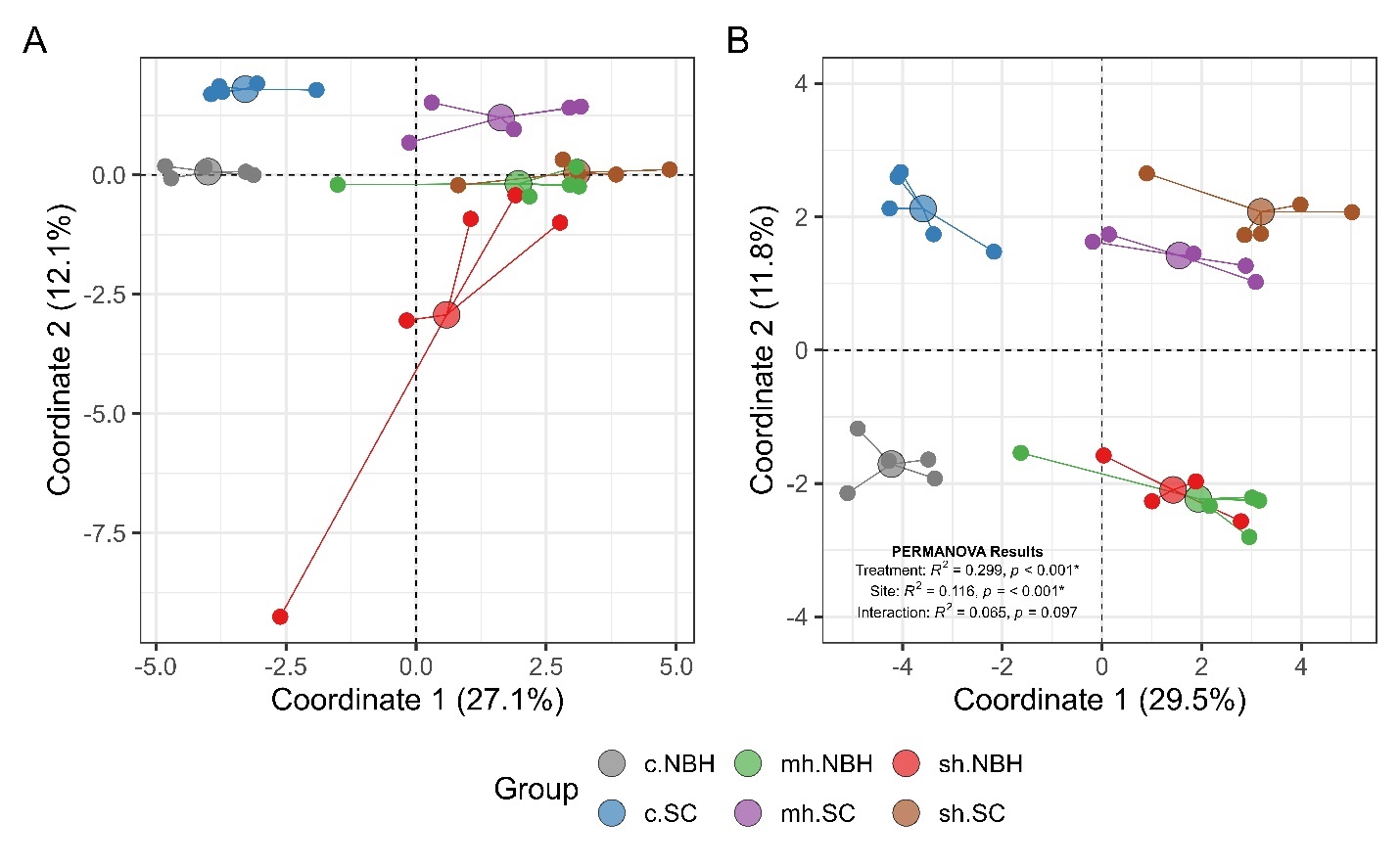

**Figure S1:** Results from the principal coordinate analysis (logCPM of 12,043 expressed transcripts) are presented for (A) all samples and (B) after outlier removal. Small colored circles represent the coordinates of individual samples (N = 5) and are connected to large colored circles, which represent the average coordinate position for each treatment group (c: control, mh: mild hypoxia, sh: severe hypoxia; SC: Scorton Creek, NBH: New Beford Harbor). The distances between points reflect the similarity in gene expression profiles among individuals and treatment groups. One NBH individual in the severe hypoxia treatment was identified as a substantial outlier and was removed from subsequent analysis. A permutational multivariate ANOVA (PERMANOVA) tested the effects of treatment and populations on global gene expression patterns (logCPM) using the R package Apev5.8, revealing significant effects of treatment and site, but not their interaction.

**Figure S2.**

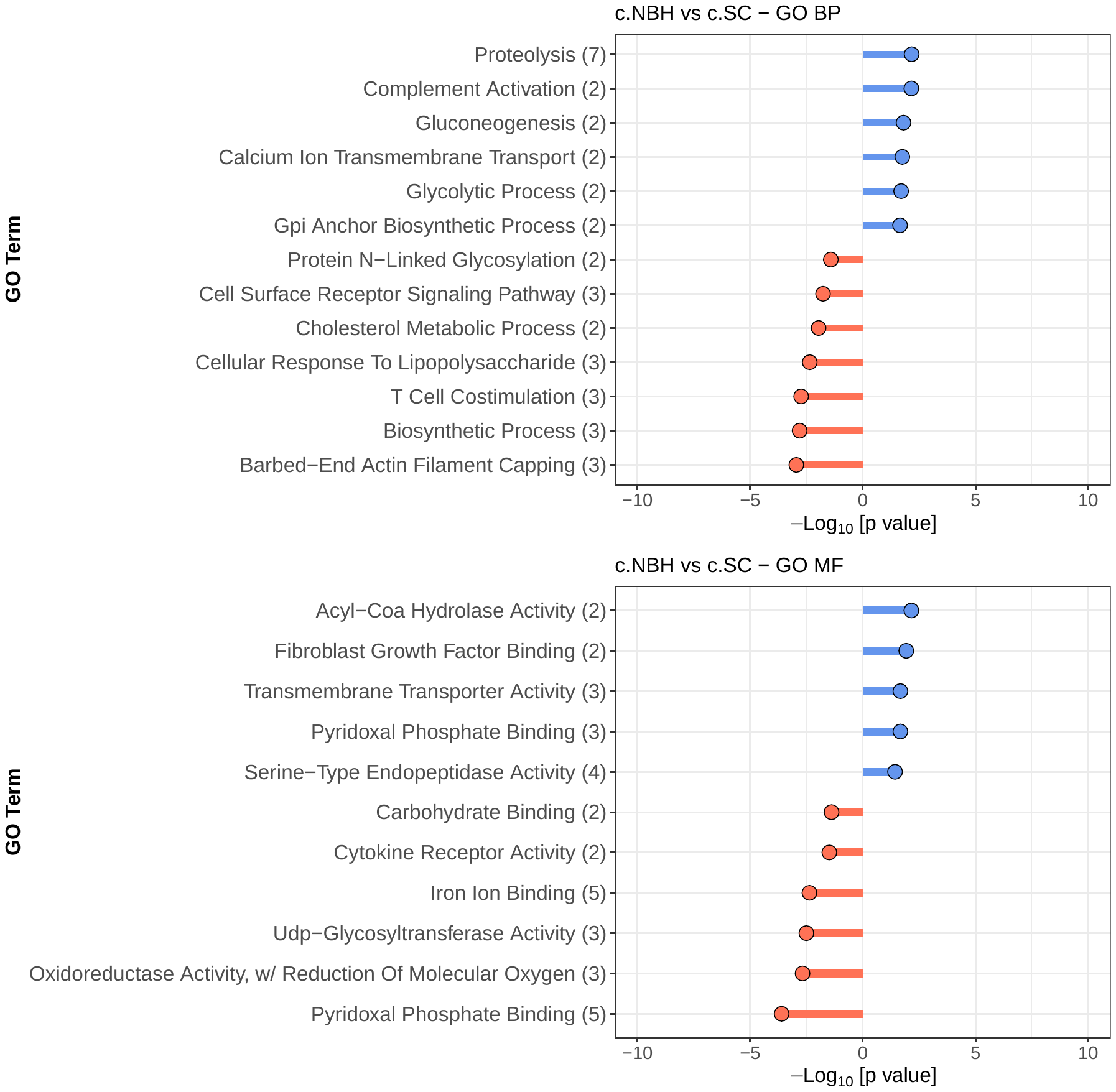
A.

B**.**

**Figure S2.** Gene Ontology terms enriched among differentially expressed genes (DEGs) between populations. Control groups between NBH and SC populations were compared. DEGs were determined using SC as reference. Only top 5 GO BP terms (A) and GO MF terms (B) are shown. Entire list of GO biological process and molecular function terms are provided in the supplementary information (RNAseq_Supplementary Information.xlsx). Detailed description of filtering of GO terms to remove redundancy is described in the materials and methods section. GO terms enriched among upregulated DEGs are in blue and those from downregulated genes are in red.

**Figure S3.**

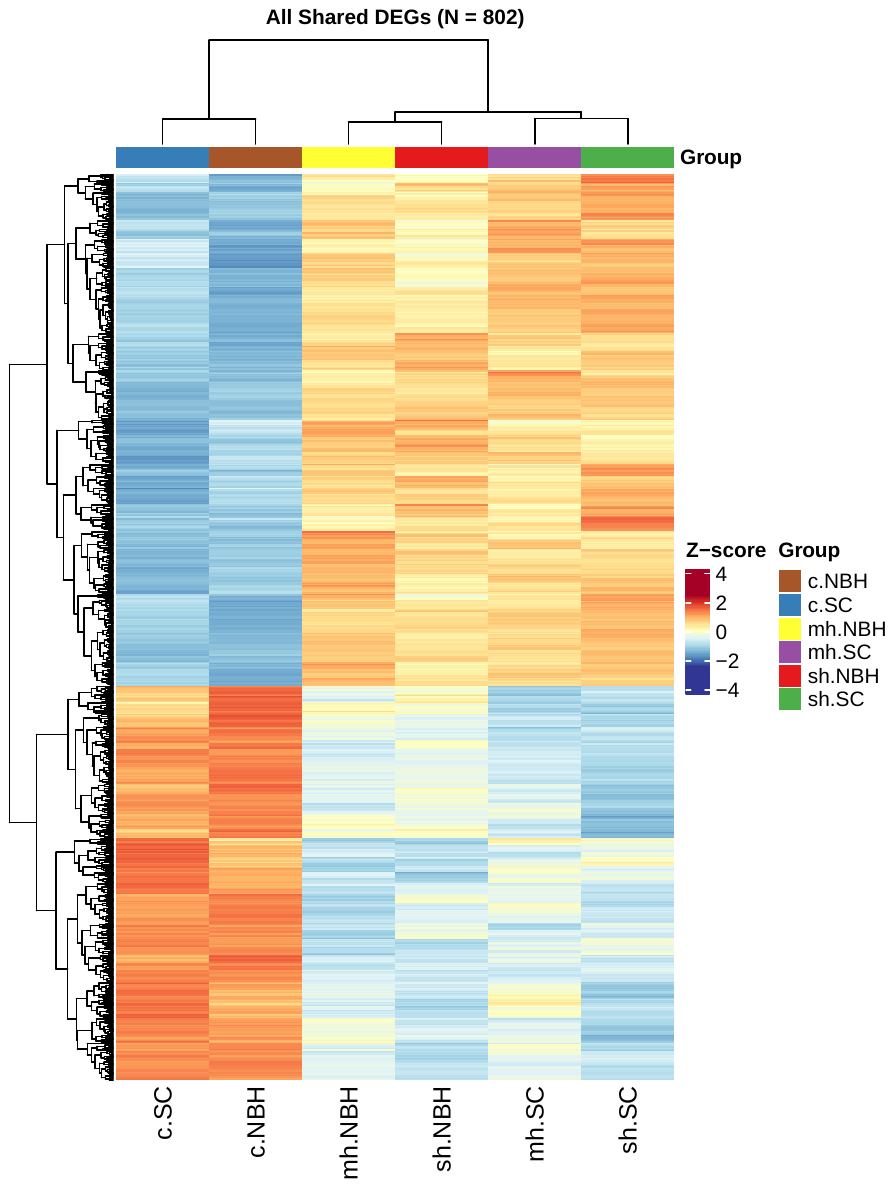

**Figure S3. Heat map representation of DEGs from all treatment comparisons.** A total of 453 upregulated and 349 downregulated genes were shared between the two populations and two hypoxia treatment groups. The expression levels (log CPM) of DEGs from each treatment are plotted. Expression level of each treatment group is the mean value from all the biological replicates. Scorton Creek (SC), New Bedford Harbor (NBH), Control (C), Mild hypoxia (mh), Severe hypoxia (sh).

**Figure S4.**

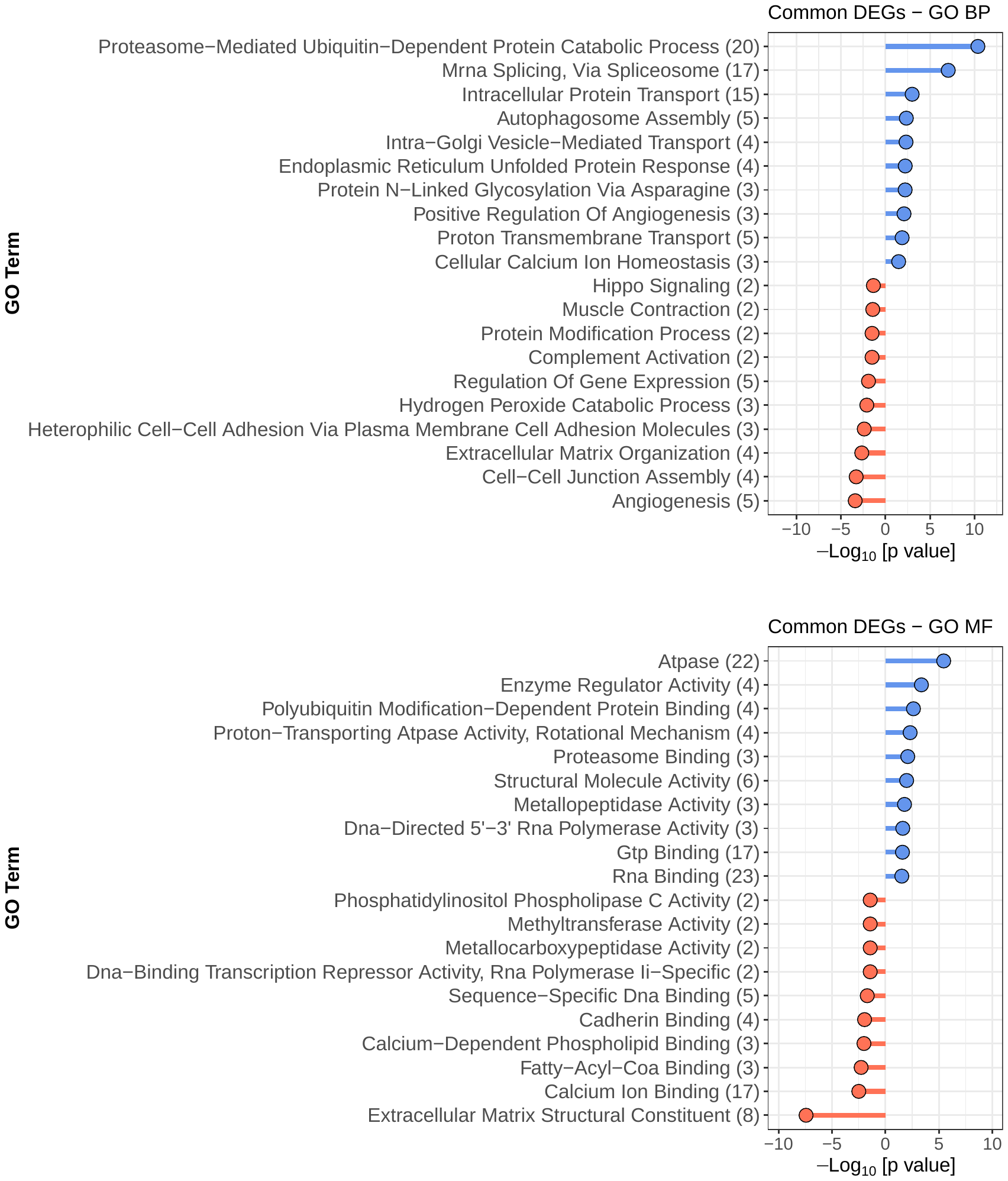

A.

B.

**Figure S4.** Gene Ontology terms enriched among differentially expressed genes (DEGs) across all treatment groups. Only top 10 GO BP terms (A) and GO MF terms (B) are shown. Entire list of GO biological process and molecular function terms are provided in the supplementary information (RNAseq_Supplementary Information.xlsx). Detailed description of filtering of GO terms to remove redundancy is described in the materials and methods section. GO terms enriched among upregulated DEGs are in blue and those from downregulated genes are in red.

**Figure S5.**

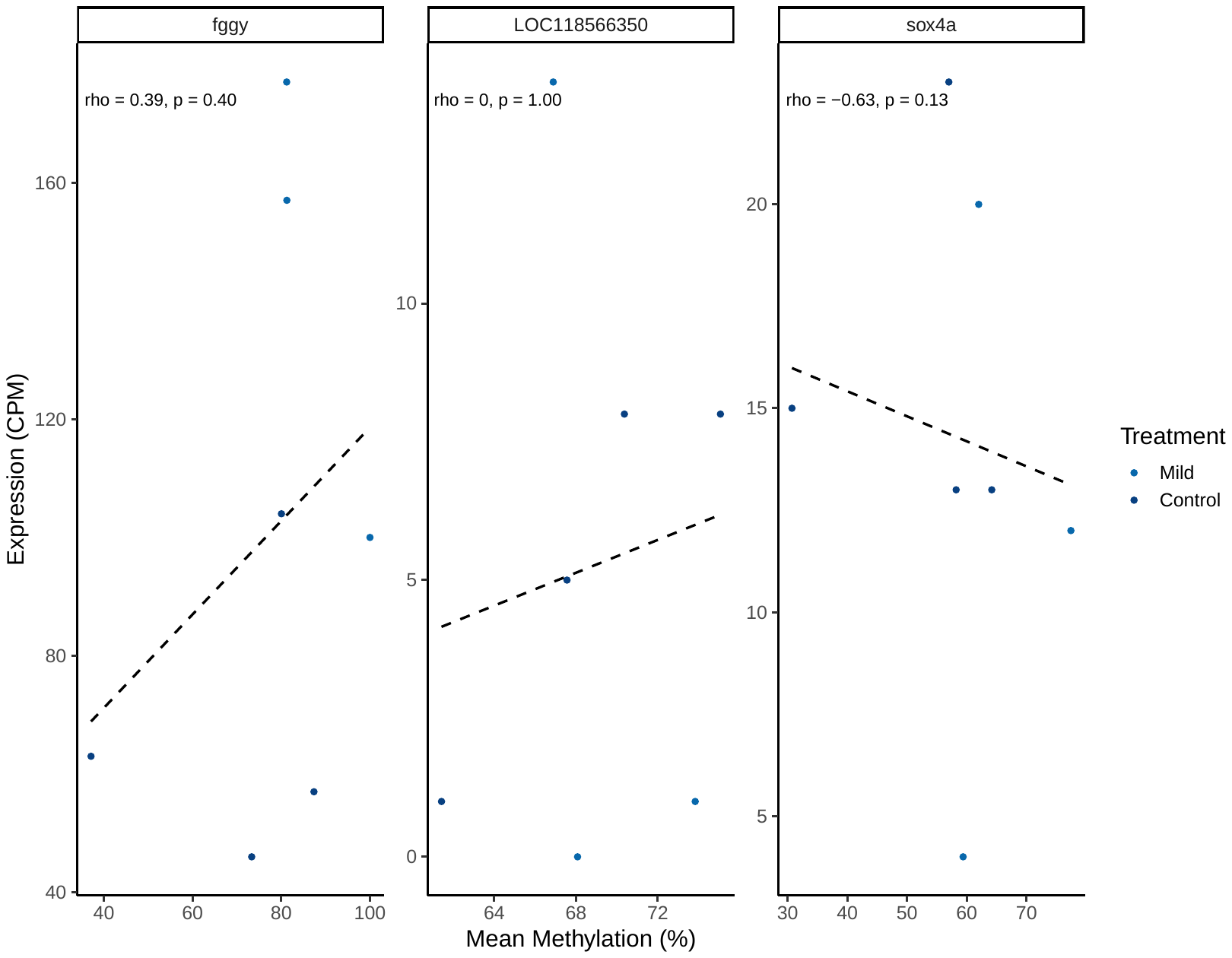

**Figure S5.** Correlation plots showing the relationship between the methylation level in DMRs and expression level of the associated gene in mild hypoxia group in New Bedford Harbor (NBH). Percent mean methylation level (x-axis) is plotted against gene expression (counts per million, CPM; y-axis). This analysis was done using BAT_correlating function in Bisulfite Anaysis Tool (BAT). Out of 10 DMRs identified in severe hypoxia NBH fish, only 3 DMRs are associated with annotated genes. We did not observe any significant correlation between mean methylation level in DMRs and the gene expression patterns.

**Figure S6.**

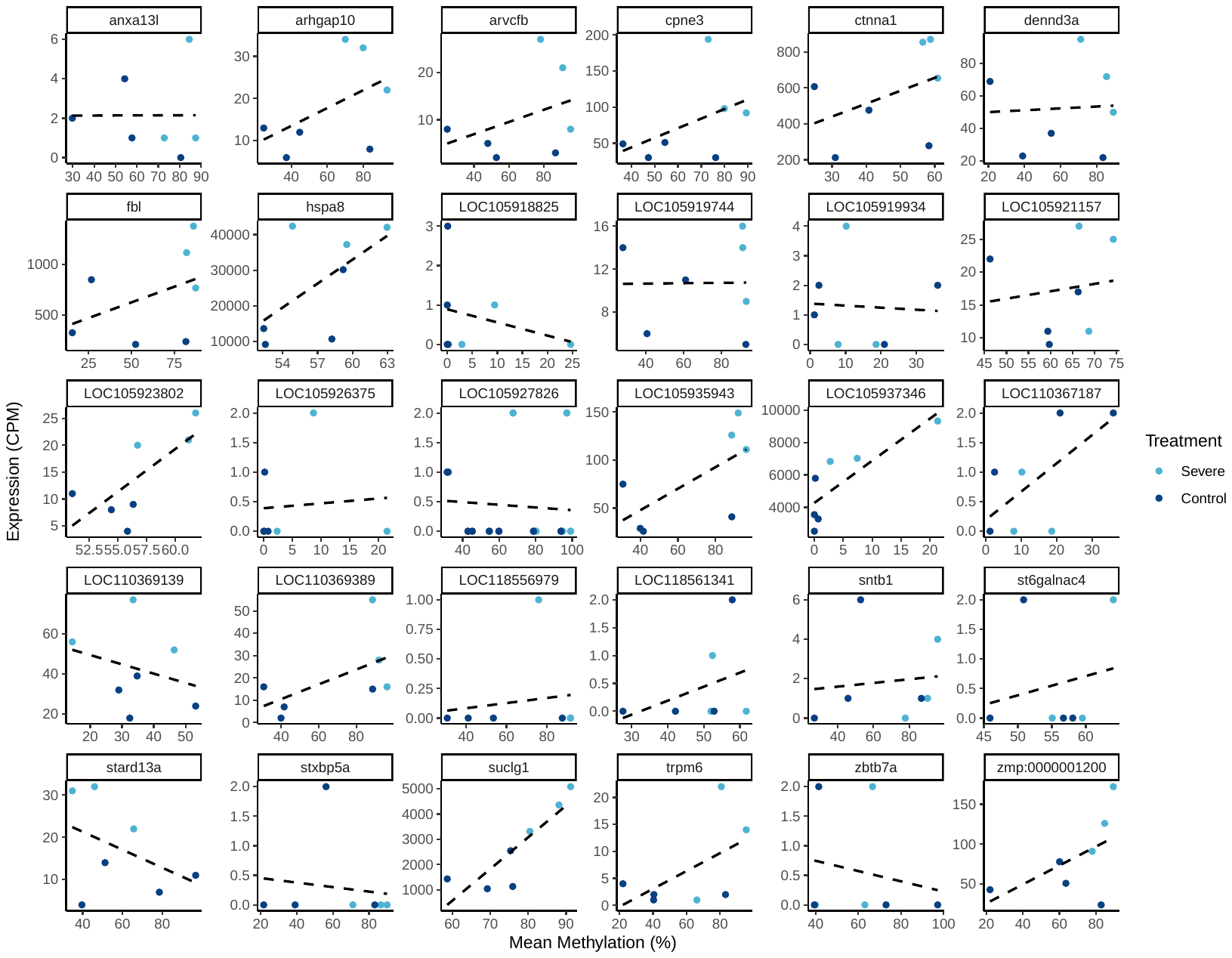

**Figure S6.** Correlation plots showing the relationship between the methylation level in DMRs and expression level of the associated gene in severe hypoxia group in New Bedford Harbor (NBH). Percent mean methylation level (x-axis) is plotted against gene expression (counts per million, CPM; y-axis). This analysis was done using BAT_correlating function in Bisulfite Anaysis Tool (BAT). Out of 59 DMRs identified in severe hypoxia NBH fish, only 30 DMRs are associated with annotated genes. We did not observe any significant correlation between mean methylation level in DMRs and the gene expression patterns.
